# Synchronous brain dynamics establish brief states of communality in distant neuronal populations

**DOI:** 10.1101/2021.01.07.425731

**Authors:** Martin Seeber, Christoph M. Michel

## Abstract

Intrinsic brain dynamics co-fluctuate between distant regions in an organized manner during rest, establishing large-scale functional networks. We investigate these brain dynamics on a millisecond time scale by focusing on Electroencephalographic (EEG) source analyses. While synchrony is thought of as a neuronal mechanism grouping distant neuronal populations into assemblies, the relevance of simultaneous zero-lag synchronization between brain areas in humans remains largely unexplored. This negligence is due to the confound of volume conduction, leading inherently to temporal dependencies of source estimates derived from scalp EEG (and Magnetoencephalography, MEG), referred to as spatial leakage. Here, we focus on the analyses of simultaneous, i.e., quasi zero-lag related, synchronization that cannot be explained by spatial leakage phenomenon. In eighteen subjects during rest with eyes closed, we provide evidence that first, simultaneous synchronization is present between distant brain areas and second, that this long-range synchronization is occurring in brief epochs, i.e., 54-80 milliseconds. Simultaneous synchronization might signify the functional convergence of remote neuronal populations. Given the simultaneity of distant regions, these synchronization patterns might relate to the representation and maintenance, rather than processing of information. This long-range synchronization is briefly stable, not persistently, indicating flexible spatial reconfiguration pertaining to the establishment of particular, re-occurring states. Taken together, we suggest that the balance between temporal stability and spatial flexibility of long-range, simultaneous synchronization patterns is characteristic of the dynamic coordination of large-scale functional brain networks. As such, quasi zero-phase related EEG source fluctuations are physiologically meaningful if spatial leakage is considered appropriately.

**Significance:** Synchrony is suggested as a mechanism for coordinating distant neuronal populations. Yet, simultaneous (i.e., zero-lag) synchronization between remote brain regions in humans is difficult to demonstrate, because volume conduction in EEG/MEG recordings causes spurious zero-lag relations. Here, we investigate actual zero-lag relations and systematically compare them to the residual bias due to spatial smoothness of EEG source estimates. We indeed report simultaneous synchronization between distant brain regions. These synchronization patterns manifest variably in time. We suggest that simultaneous synchronization is relevant when studying the dynamic, large-scale functional architecture in humans.

## Introduction

Brain activity spontaneously fluctuates during rest, when no specific task is instructed. Intriguingly, these fluctuations are correlated between distant brain regions, forming large-scale functional networks that are assumed to reflect spontaneous information integration during internal mentation (Raichle et al., 2001; Greicius et al., 2003; Smith et al., 2009; Brookes et al., 2011; Engel et al., 2013), i.e., the basis of thinking. While functional magnetic resonance imaging (fMRI) was crucial for the discovery and investigation of resting-state networks, the low time resolution of BOLD variations does not allow us to study the neurophysiological mechanisms leading to these spontaneous co-fluctuations of spatially distinct brain areas. Intracranial local field potential recordings or scalp electro-/magnetoencephalography (EEG/MEG) are adequate for this purpose, as they record neuronal activity at their inherent time-scale, i.e. in the millisecond range (Roelfsema et al., 1997; Miller et al., 2009; Baker et al., 2014; Fox et al., 2018; Vidaurre et al., 2018). Such studies revealed an essential key neuronal mechanism underlying information integration between different brain regions: Synchrony (Singer, 1999; Varela et al., 2001). Many studies have demonstrated that neuronal synchronization between brain areas is an important mechanism for the coordination of neuronal processing in anatomically distributed neuronal circuits (Engel et al., 1991; Contreras and Steriade, 1996; Roelfsema et al., 1997; Destexhe et al., 1999; Womelsdorf et al., 2007). A fundamental question is whether synchronous co-fluctuations between areas are simultaneous or time-lagged (Engel et al., 1991; Contreras and Steriade, 1996; Roelfsema et al., 1997; Destexhe et al., 1999; Fries, 2005; Womelsdorf et al., 2007; Siegel et al., 2008; Bosman et al., 2012; Van Kerkoerle et al., 2014). Because of delays due to axonal conduction and synaptic transmission, time-lagged fluctuations are necessarily appearing when the activation of one region is causally related to the activation of the other region, i.e. when one area transfers information to the other. Simultaneity, on the other hand, indicates a gathering of different brain areas converging into a functional unit to collectively maintain certain information without causal interactions between them. Such communality can be established spontaneously by dynamic recurrent connections or can be driven by a pacemaker (e.g., the thalamus) (Vicente et al., 2008; Gollo et al., 2014). Undoubtedly, both mechanisms (time-lagged and simultaneous fluctuations) take place in the brain to processes, integrate and maintain the information, as numerous intracranial recordings in animals and humans have shown (Contreras and Steriade, 1996; Roelfsema et al., 1997; Womelsdorf et al., 2007; Siegel et al., 2008; Hipp et al., 2011). Unfortunately, simultaneous activity, which imposes zero-lag related signals are primarily ignored in EEG/MEG network analyses to avoid spurious phase relations resulting from volume conduction (Nolte et al., 2004; Stam et al., 2007; Hipp et al., 2012; Marzetti et al., 2013; Colclough et al., 2015). EEG/MEG source reconstruction (Michel et al., 2004; Michel and Murray, 2012; He et al., 2018) is, to some extent, able to overturn volume conduction effects. Yet, the limited spatial resolution of EEG/MEG source reconstruction techniques leads to spurious temporal relations (Palva et al., 2018; He et al., 2019). To correct for these spatial leakage effects, orthogonalization of source signals is a standard method. However, this method also discards genuine simultaneous dynamics and therefore is insensitive to detect such.

In this work, we aim to investigate simultaneous synchronization, i.e., quasi zero-lag relations between distant brain areas using high-density EEG source imaging (Michel et al., 2004; Michel and Murray, 2012; He et al., 2018). To consider and correct for spatial leakage effects, we systematically compare actual with surrogate data having identical spatial properties in their source reconstruction.

In summary, we demonstrate that physiologically meaningful quasi zero-lag synchrony between distant brain areas exists that cannot be explained by spatial leakage phenomena. We suggest that brief epochs of simultaneous synchronization signify functional convergence of distant neuronal population dynamics into distinct re-occurring states.

## Methods

### EEG recordings

High-density EEG was recorded using an electrode net (Geodesic Sensor Net, Electrical Geodesics Inc., Eugene, OR, USA) consisting of 256 electrodes that are interconnected by thin rubber bands. Each electrode includes a small sponge soaked with saline water to establish direct electrical contact with the participants’ scalp. EEG was sampled at 1 kHz, referenced to the vertex.

Participants (N=18, 30 ± 5 years, seven male) sat comfortably in an upright position in a darkened, electrically shielded room and were instructed to keep their eyes closed and relax for four to six (5.42 ± 0.95) minutes avoiding drowsiness. The local ethical committee, following the declaration of Helsinki, approved the study. Participants provided written, informed consent for their participation.

### EEG preprocessing

EEG recordings were band-pass filtered between 1-40 Hz offline, and electrodes covering cheeks and nape were excluded. Time epochs contaminated with apparent artifacts were marked and excluded from further analyses. Noisy or bad electrodes were excluded from Independent Component Analysis (ICA) (Jung et al., 2000), which was used to remove stereotypical artifact components containing saccades, eye blinks, and cardiac artifacts. Afterward, the initially excluded channels were spline interpolated in space, resulting in 204 channels. The recordings were re-referenced to the common average and down-sampled to 125 Hz for further analysis.

### EEG source imaging and functional network reconstruction

We applied EEG source reconstruction using forward models based on realistic head geometry and conductivity data with consideration of skull thickness, i.e., Locally Spherical Model with Anatomical Constraints (LSMAC) (Brunet et al., 2011; Michel and Brunet, 2019). The grey matter was defined based on the MNI anatomical template model. The inverse solution space consisted of 5004 points equally distributed in this grey matter volume. The linear distributed inverse solution LAURA (Grave de Peralta Menendez et al., 2004) was used to calculate the current density distribution for each solution point at each moment in time. Dipole orientations were set to the first left singular vector of the xyz (3D) components in the resolution matrix of each source pointing outside of the brain to avoid sign ambiguities.

Functional networks were defined as spatial patterns co-varying with fluctuations in selected regions of interest (ROI) defined in an atlas parcellation (Schaefer et al., 2017). We chose the posterior cingulate cortex (PCC) and the supplementary motor area (SMA) as two exemplary seed regions based on previous literature focusing on functionally distinct key regions (Seeley et al., 2007; Raichle, 2010; Engel et al., 2013). The signal representing the activities in each ROI was defined as the first principal component of all dipoles within the given ROI (Rubega et al., 2018). Then, we calculated their signal envelope as the magnitude of the analytic signal using the Hilbert transform. To capture well-pronounced spatial patterns that include these key regions, we thresholded the signal envelope at the mean plus standard deviation following previous work (Tagliazucchi et al., 2012). The network patterns were then determined by sites that covary with this seed signal. To illustrate the resulting spatial patterns, they were spatially thresholded using watershed transform, and the local maxima positively co-varying with the respective ROI are shown (Fig. 1).

**Figure 1.**
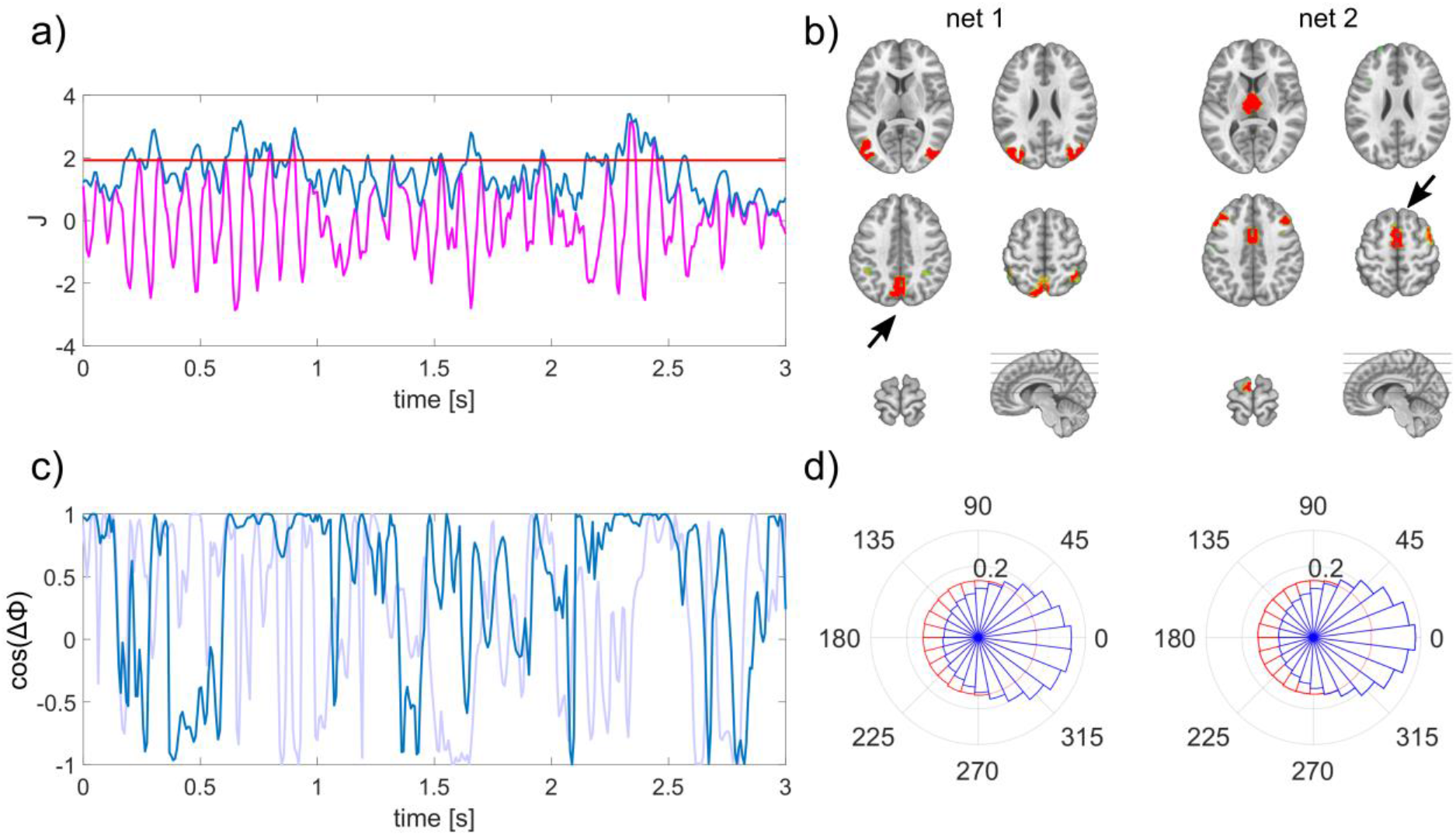
Derivation and characterization of EEG source reconstructed networks. a) The envelope (blue) of source estimated activity (magenta) is thresholded to define periods of well-pronounced activity within a specific region of interest (here PCC). b) Nodes of the network co-varying with the PCC (net 1) during periods defined as indicated in a) and with the SMA (net 2) as region of interest marked with black arrows. c) Exemplary time course of instantaneous phase locking between lateral posterior regions of net 1, matching the time period shown in a) in magenta; surrogate phase locking is shown in light blue. d) Polar histograms of the group, displaying the distribution of interhemispheric phase differences between lateral posterior (net 1) and anterior (net 2) regions as illustrated in b) in blue; surrogate phase differences in red. The radius for each phase bin displays the probability density function estimate of the respective phase differences.

### Surrogate data and spatial leakage estimation

To systematically asses the bias introduced by spatial leakage we used surrogate data, which we derived from the actual data. To do so, we temporally shifted the source reconstructed signals of the actual data randomly in time for every solution point individually for each subject. That way, the initial source dynamics of the surrogate data are the same as the actual source estimates, but the temporal relations between solution points are demolished. To introduce spatial leakage, we then applied the same forward model as used for analyzing actual data to generate surrogate EEG. Afterwards, we applied the identical processing pipeline to this surrogate data, i.e. filtering scalp data and source estimation using the same inversion kernel as in the analyses of the actual EEG data. Because we used identical forward model and inverse method for analyzing actual and surrogated data, the spatial properties of the source estimates are the same. That way, there are no actual correlations between the sources given the introduced random time shifts. Therefore, the resulting inter-areal correlation values in the surrogate source estimates are due to spatial leakage between selected areas. This procedure provides bias estimates caused by spatial leakage for every connectivity metric, i.e. correlation, phase-locking value (PLV) and coherence for each individual subject. These bias estimates can be subtracted from the metrics of actual data as suggested previously (Ghuman et al., 2011; Palva and Palva, 2012) and used for statistical comparison.

### Synchrony between network nodes

We investigated the correlation, lag, phase locking and coherence between network nodes. Between each pair, we determined the correlation for different lags of the signals using cross-correlation. To perform frequency-specific analyses, we applied wavelet transform (Morlet et al., 1982) for time-frequency (TF) decomposition (1–40 Hz, 1 Hz steps). Parameters for the mother wavelet were set to the full width at half maximum of three seconds for the Gaussian kernel at a center frequency of 1 Hz. PLV and coherence was computed for every frequency bin and are reported as magnitudes herein and for the latter as real and imaginary part of the coherency (Lachaux et al., 1999; Lachaux et al., 2002) to compare with previous literature (Nolte et al., 2004). Simultaneous synchrony is indicated as peak correlation at zero-lag in the cross-correlogram and the real part of coherency. The time-varying phase in each ROI was computed using Hilbert transform in order to determine phase differences between regions for every time point. The distribution of these phase differences were illustrated as polar histograms. The cosine of these phase differences ∆φ was used as instantaneous measure of simultaneous synchronization, which is 1 for zero phase difference (Deco and Kringelbach, 2016; Cabral et al., 2017). The duration of phase synchrony, which is centered around zero phase lag was determined by epochs of cos(∆φ) exceeding 0.5. Very short epochs smaller than 24ms, i.e. 3 time samples, were not considered as stable and therefore ignored for computing the average duration. All metrics were statistically compared to results derived from surrogate data. Paired comparisons were carried out using the Wilcoxon signed-rank test, which were Bonferroni corrected for multiple comparisons.

## Results

### Large-scale brain dynamics form briefly stable functional networks

We found bilateral, symmetric posterior regions in the extrastriate cortex and inferior parietal lobe (IPL) to co-vary with the PCC’s source signal. In contrast, we found anterior areas of the bilateral prefrontal cortex and the thalamus to co-vary with the SMA (Fig. 1a-b). To rule out a potential source imaging bias that might cause these patterns, we performed the same analyses on the surrogate data. Importantly, we found no distant spatial local maxima forming a network pattern in the surrogate data. Merely the respectively selected regions were present, meaning we did not observe co-varying regions using surrogate data (Fig. 6).

The phase relations between nodes of these functional network patterns in the real data vary considerably in time. We observe epochs in which the phase differences remain small, meaning these two nodes fluctuate synchronously at these time points (Fig. 1c-d). The durations of these epochs are in the range between 54.1 and 79.1 milliseconds on average depending on the constellation. The durations of all pairs belonging to the same functional network are significantly longer than respective periods computed from surrogate data. The detailed duration of each pair and their respective p-values are listed in Table 1.

**Table 1.** Correlation, PLV in the alpha range (8-12 Hz) and duration of each pair with respective p-values (Wilcoxon sign rank test, Bonferroni corrected)

|  | r | p<sub>r</sub> | PLV | p<sub>PLV</sub> | Dur [ms] | p<sub>Dur</sub> |
| --- | --- | --- | --- | --- | --- | --- |
| left IPL - PCC | 0.20 | 0.0038 | 0.22 | 0.0011 | 69.9 | 0.0007 |
| left IPL - right IPL | 0.28 | 0.0007 | 0.28 | 0.0007 | 79.1 | 0.0007 |
| right IPL - PCC | 0.10 | 0.0123 | 0.15 | 0.0007 | 66.2 | 0.0024 |
| left PFC- SMA | 0.24 | 0.0012 | 0.34 | 0.0007 | 57.1 | 0.0038 |
| left PFC - right PFC | 0.31 | 0.0007 | 0.37 | 0.0007 | 54.1 | 0.0020 |
| right PFC – SMA | 0.26 | 0.0012 | 0.34 | 0.0007 | 61.9 | 0.0009 |
| SMA – thalamus | 0.32 | 0.0020 | 0.34 | 0.0012 | 75.8 | 0.0011 |

### Simultaneous synchronization is present between distant neuronal populations

We identified functional network patterns that are composed of distinct nodes that are symmetric in both hemispheres (Fig. 1). This finding already indicates that these distant regions co-vary on a highly resolved time scale. To directly test if the correlation between these nodes is significantly larger than the spatial leakage bias, we focused on the analyses of pairwise nodes for each network pattern. To provide more detail about these interactions, we investigated different time lags and frequency components. For the PCC based network, we focused on posterior bilateral IPL regions. The cross-correlation between pairs of these network nodes peaks at zero-lag with values ranging between 0.1 and 0.28, which is significantly higher than the spatial leakage bias observed in the surrogate data. The detailed values are listed in Table1. Interestingly, the interhemispheric zero-lag correlation was highest in this posterior network. The frequency-specific PLV reached its maximum for this pair at 11Hz with a value of 0.34. In this case, the real part of the coherency is considerably higher than its imaginary part (Fig.2).

**Figure 2.**
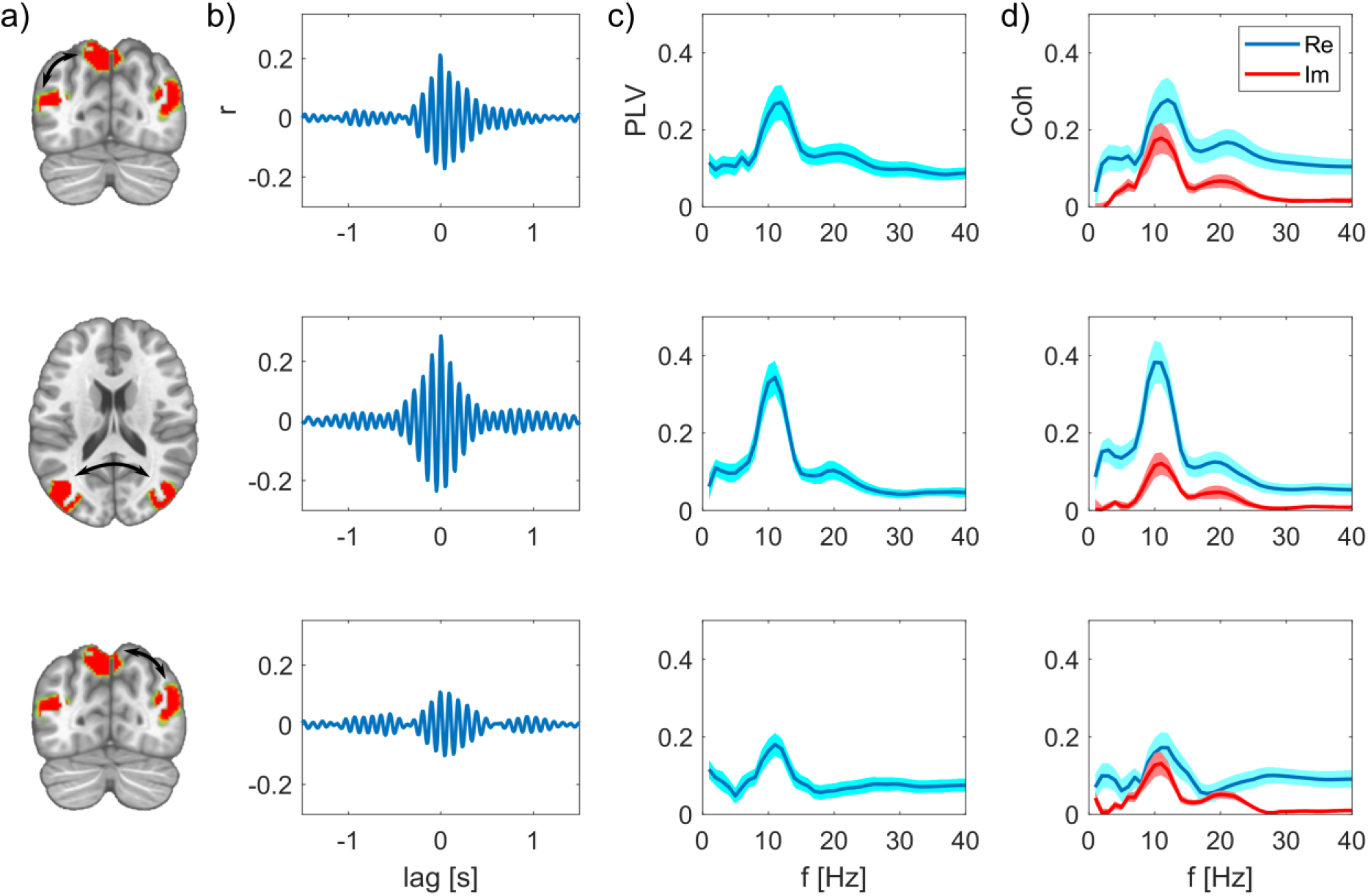
Synchrony between the nodes of the PPC network after subtracting spatial leakage bias. a) Nodes of the network, edges are indicated as arrows. b) Cross-correlations between these two nodes are respectively maximal at zero lag. c) PLV as function of frequency, group mean ± SEM. d) Real and imaginary part of the coherency, group mean ± SEM.

For the SMA based network, we further examined the relation of the SMA to regions in the bilateral PFC and to the thalamus. The cross-correlation between these regions peaks at zero-lag with a value of ranging between 0.24 and 0.32, which is significantly higher than the spatial leakage bias observed in the surrogate data. The frequency-specific PLV reached its maximum at 10Hz with a value of 0.42 for the interhemispheric PFC connection. Again, the real part of the coherency is higher than its imaginary part (Fig.3). These results show that actual zero-phase relations, indicating simultaneous synchronization, are present between relatively distant regions.

**Figure 3.**
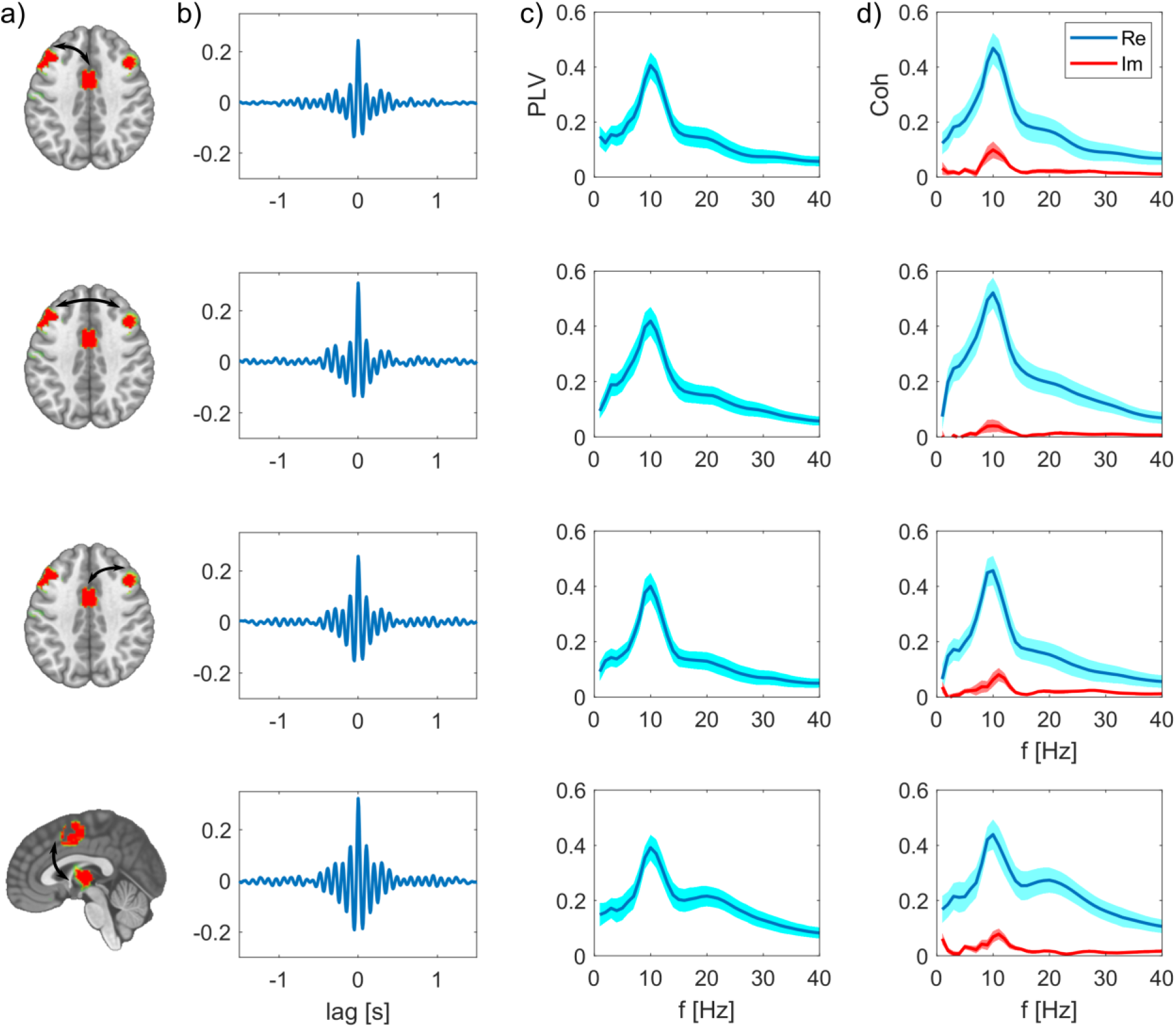
Synchrony between the nodes of the SMA network after subtracting spatial leakage bias. a) Nodes of the network, edges are indicated as arrows. b) Cross-correlations between these two nodes are respectively maximal at zero lag. c) PLV as function of frequency, group mean ± SEM. d) Real and imaginary part of the coherency, group mean ± SEM.

For direct visual comparison of actual with surrogate data we also show the uncorrected metrics overlaid with the bias estimates in Fig.4 and Fig.5. These bias estimates are the higher, the closer a node pair is, but also the lower the spatial resolution between these areas is. For example, the zero-lag correlation peak of the surrogate data is higher for the intrahemispheric pairs (Fig.4b, top and bottom row), than the bias of the more distant interhemispheric pair (Fig.4b middle row). This is analogously the case for the PLV relations in Fig.4c. The same applies for comparing the top three rows in Fig.5c-d for the SMA based network. The bias due to spatial leakage is maximal between SMA and the thalamus, which is plausible given the low spatial resolution in subcortical areas (Fig.5, bottom row). In addition, spatial leakage is biasing the phase distribution of the surrogate data towards zero, i.e. right in the plots of Fig.4d and Fig.5d. In other terms, the phase distribution is not circular any more, but biased due to spatial leakage, which is best visible in Fig.5d, bottom row (displayed in red). Yet, for the actual recordings, the phase bin centered around zero exceeds this bias significantly (displayed in blue).

**Figure 4.**
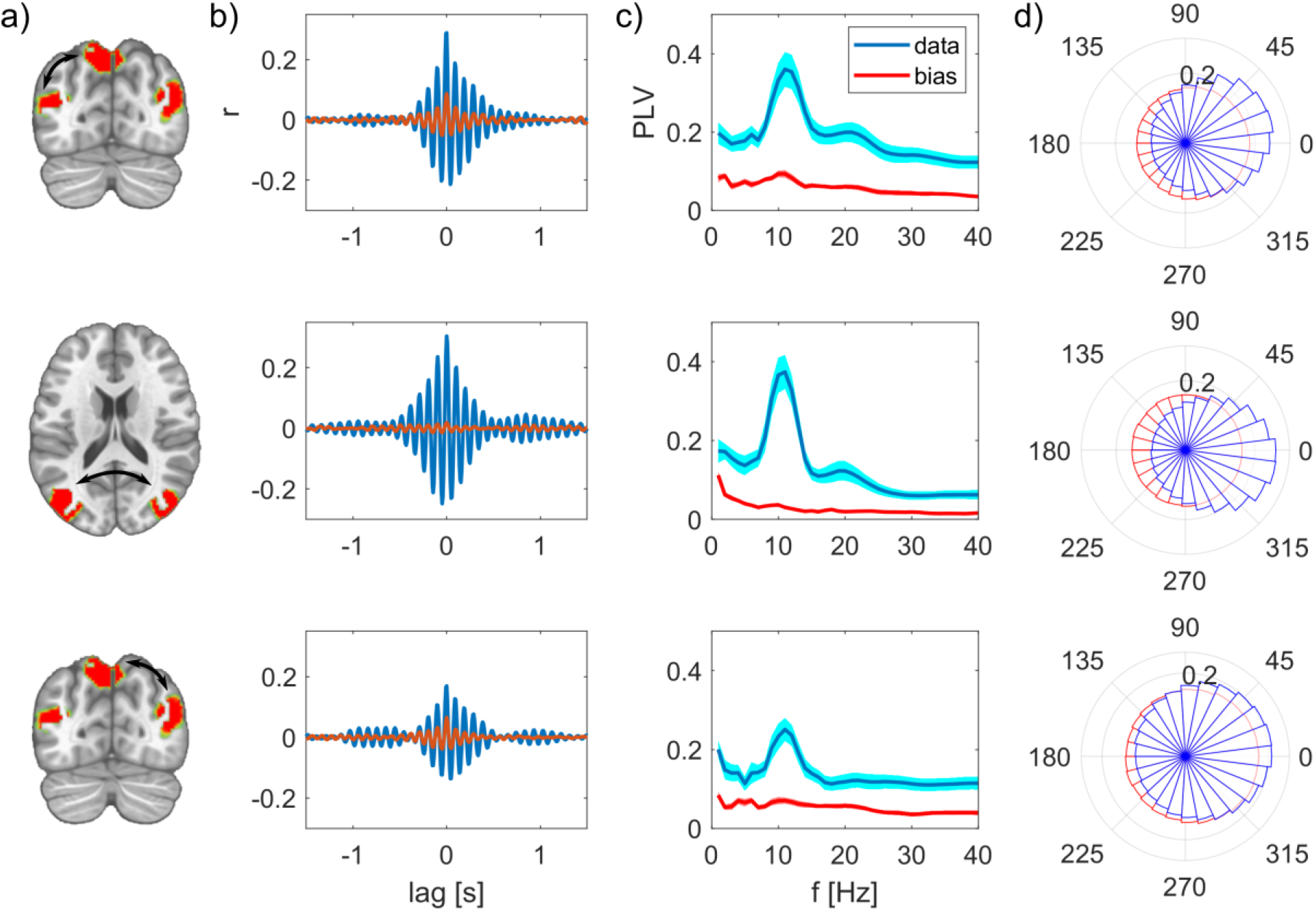
Synchrony between the nodes of the PPC network, uncorrected measures in comparison to spatial leakage bias. a) Nodes of the network, edges are indicated as arrows. b) Cross-correlation between these two nodes, actual (uncorrected) data is shown in magenta, bias in surrogate data in red. c) PLV as function of frequency, group mean ± SEM. d) Polar histograms showing the distribution of phase differences for actual data in blue and surrogate data in red. The radius for each phase bin displays the probability density function estimate of the respective phase differences.

**Figure 5.**
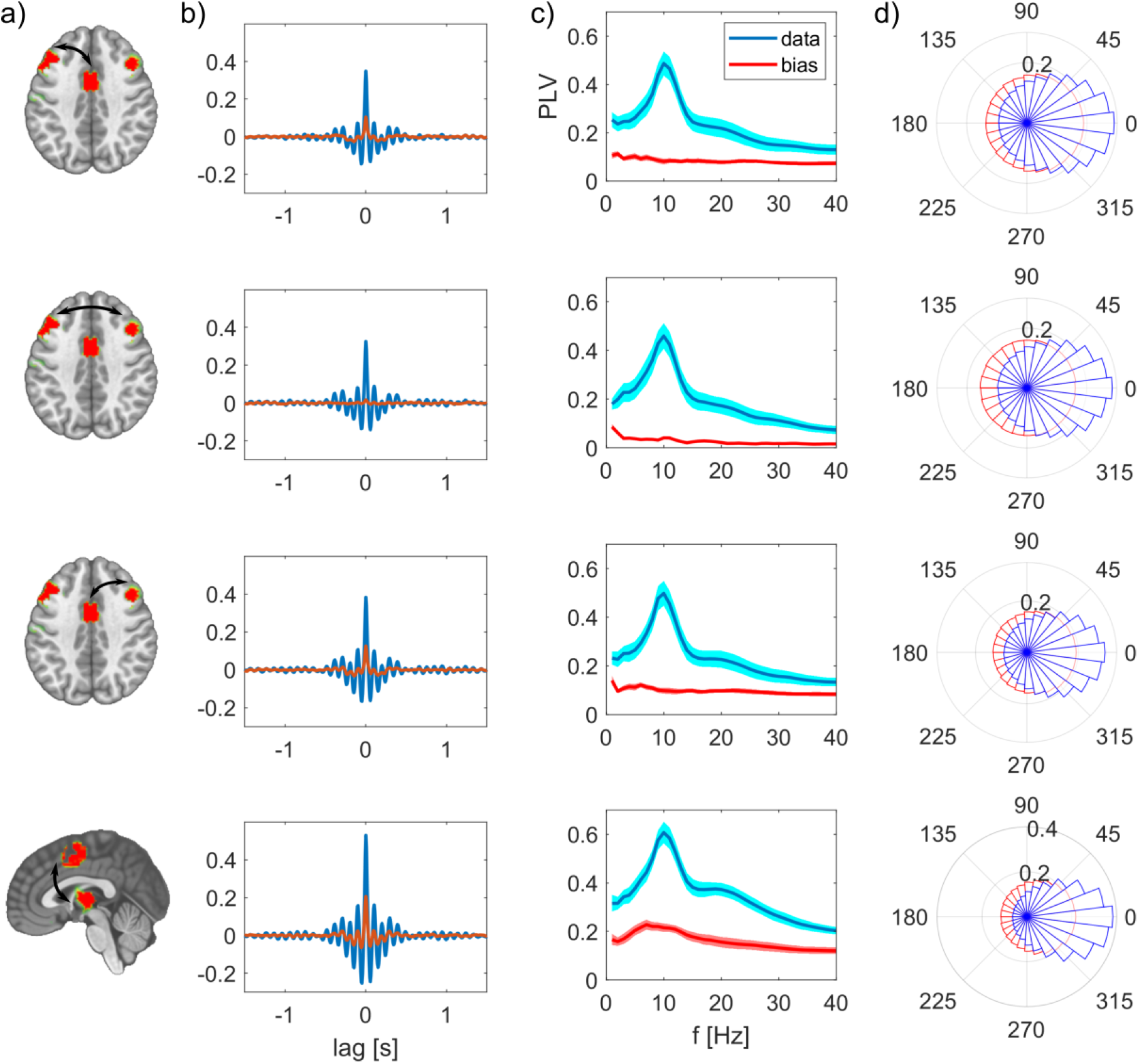
Synchrony between the nodes of the SMA network, uncorrected measures in comparison to spatial leakage bias. a) Nodes of the network, edges are indicated as arrows. b) Cross-correlation between these two nodes, actual (uncorrected) data is shown in magenta, bias in surrogate data in red. c) PLV as function of frequency, group mean ± SEM. d) Polar histograms showing the distribution of phase differences for actual data in blue and surrogate data in red. For every constellation in this network, the most frequent phase difference is zero. The radius for each phase bin displays the probability density function estimate of the respective phase differences.

## Discussion

In this work, we investigate synchronous EEG source dynamics between distant brain regions. The functional network patterns we reconstruct revealed spatially well-separated remote brain regions. Investigating the temporally highly resolved phase relations indicating long-range synchronization, we actually observe quasi zero-lag related fluctuations between these distant regions. By comparing these results systematically to surrogate data with identical spatial properties in their source reconstruction, we demonstrate that the observed effects cannot be explained by spatial leakage phenomena.

### Large-scale brain dynamics form briefly stable functional networks

In the reconstruction of functional network patterns, we focused on two key brain regions, i.e., the PCC and SMA. The PCC based network is composed of bilateral posterior areas of the extrastriate cortex and inferior parietal lobes. This network resembles the posterior subdivision of the default mode network that was previously reported using MEG recordings (Hipp et al., 2012; Vidaurre et al., 2018). The SMA based network is composed of the bilateral prefrontal cortex and the thalamus, which are regions associated with the anterior part of the control network (Seeley et al., 2007; Raichle, 2010). We included analyses of thalamic signals because recent work (Krishnaswamy et al., 2017; Seeber et al., 2019) demonstrated the detectability of subcortical activities using EEG source imaging.

However, we did not find a one to one correspondence between the network patterns we observed herein and the M/EEG amplitude correlation-based networks (Brookes et al., 2011; Samogin et al., 2019) that were related to the well-known fMRI resting-state networks (Smith et al., 2009; Raichle, 2010). This discrepancy might stem from the different time-scale of co-variation, i.e., the temporal precision, and coupling measure, which define these functional networks. In this work, phase relations are relevant, since we were aiming for high temporal precision reflecting long-range synchrony. In contrast, in fMRI and M/EEG amplitude envelope-based analyses, the temporal alignment on a second scale is sufficient for capturing correlated activities. Phase coherence and amplitude envelop correlation are two types of coupling measures suggested to reflect distinct mechanisms related to different functions (Engel et al., 2013).

We report the nodes of these network patterns synchronizing in brief time intervals, typically in the range of 54 and 80 milliseconds. These briefly stable epochs and their duration are in good agreement with previously reported time epochs for the EEG microstates (Michel and Koenig, 2018) and transient states derived from MEG recordings using Hidden Markov Models (HMM) (Vidaurre et al., 2018). However, the HMM states are derived from orthogonalized signals (Colclough et al., 2015) that discards zero-phase relations. EEG microstates are defined as stable topographies. If a particular source network configuration maintains quasi-zero phase relations for a certain period, that necessarily leads to a stable topography of the scalp potential field. Therefore, the brief manifestation of specific quasi zero-lag related network patterns we describe in this work can be seen as the underlying source dynamics of the microstates.

The temporal dynamics of these briefly stable epochs are characteristic for metastability, i.e., signified by a counterbalance between integrated, i.e. synchronous, and segregated epochs (Tognoli and Kelso, 2014; Deco et al., 2015). In terms of large-scale brain dynamics that means specific nodes of a network pattern are converging into synchrony, i.e. quasi zero-lag relationships, for brief epochs. These integrated, highly synchronous states fall abruptly apart, i.e. segregate, before the next integrated state is established. In that way, it is possible to develop dynamic representations flexibly since distinct states can be installed in different spatial configurations (Tononi and Edelman, 1998; Deco and Kringelbach, 2016; Ju and Bassett, 2020).

### Simultaneous synchronization is present between distant neuronal populations

The fact that we observe spatially well-separated, co-varying sites as network patterns is the first indicator that these distant regions are functionally related at a millisecond time scale. These distant sites are absent when repeating these analyses with surrogate data (Fig.6). In addition to this spatial assessment, the functional results, e.g. PLVs, we describe herein significantly exceed the bias due to spatial leakage, which we derive from surrogate data. As expected, these bias estimates are the higher, the closer two areas are and the lower the spatial resolution at these sites is. Surprisingly, we found the interhemispheric interactions to be higher than the intrahemispheric interactions. Because the distance between respective regions is larger for the interhemispheric than the intrahemispheric pairs, this result cannot be an effect of spatial leakage. These findings together with the cross-correlation peak at zero lag signify genuine simultaneous synchronization between these distant regions.

**Figure 6.**
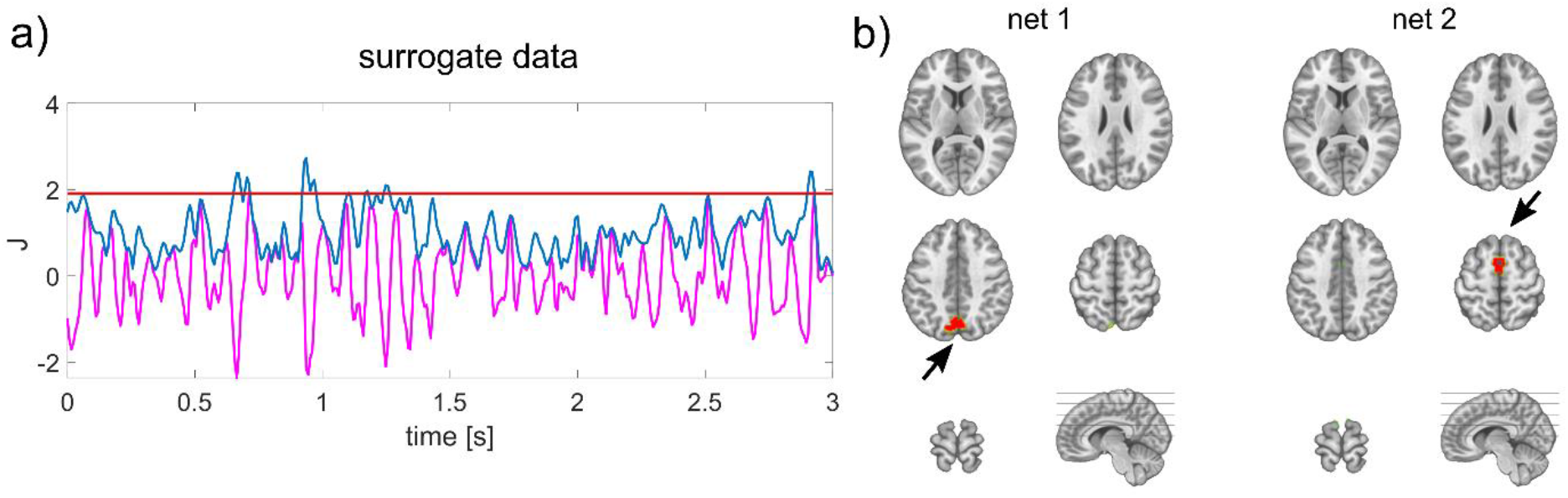
Absence of distant co-varying sites in surrogate data. a) The envelope (blue) of source estimated surrogate data (magenta) is thresholded to define periods of well-pronounced activity within a specific region of interest (here PCC). b) No distant local maxima were identified co-varying with the PCC (net 1), or with the SMA (net 2) marked with black arrows.

The finding of long-range, simultaneous synchronization is in line with previous literature showing physiologically relevant, zero-lag relations (Engel et al., 1991; Contreras and Steriade, 1996; Roelfsema et al., 1997) in animals. Recently, a study using intracranial recordings showed interhemispheric zero-lag synchronization in the human brain (O’reilly and Elsabbagh, 2020). Most of the previous studies investigating synchrony between distant areas were focusing on gamma oscillations (>30 Hz) induced by specific tasks (Engel et al., 1991; Roelfsema et al., 1997; Womelsdorf et al., 2007; Siegel et al., 2008; Van Kerkoerle et al., 2014). These gamma oscillations were found to facilitate feedforward processing, while mid-frequencies were related to feedback effects from higher areas (Von Stein et al., 2000; Bosman et al., 2012; Van Kerkoerle et al., 2014). Given these differences in task-induced and resting-state signals, it is plausible that the simultaneous fluctuations we describe here represent intrinsic synchrony during minimal sensory input. The finding of quasi zero-phase relations between distant areas might signify functional convergence in these regions during rest, in contrast to sensory-driven time-lagged oscillations induced by a specific task. In that sense, quasi zero-phase relations in distributed areas might relate to the representation and maintenance, rather than the processing of information. This long-range synchronization is briefly stable, not persistently, indicating flexible spatial reconfiguration pertaining to the establishment of particular, re-occurring states. Taken together, we suggest that the balance between temporal stability and spatial flexibility of long-range, simultaneous synchronization patterns is characteristic of the dynamic coordination of large-scale functional brain networks. As such, quasi zero-lag related EEG source fluctuations are physiologically meaningful if spatial leakage is considered appropriately, and should not be excluded in the analysis of functional connectivity using EEG/MEG source imaging.

## Acknowledgements

This study was supported by the Swiss National Science Foundation (Grant Number: 320030_184677) to C.M.M. The authors would like to thank Miralena I. Tomescu for her support in data preparation.

## Notes

### Competing Interest Statement

The authors have declared no competing interest.

## References

Baker AP, Brookes MJ, Rezek IA, Smith SM, Behrens T, Smith PJP, Woolrich M (2014) Fast transient networks in spontaneous human brain activity. Elife 3:e01867.

Bosman CA, Schoffelen J-M, Brunet N, Oostenveld R, Bastos AM, Womelsdorf T, Rubehn B, Stieglitz T, De Weerd P, Fries P (2012) Attentional stimulus selection through selective synchronization between monkey visual areas. Neuron 75:875–888.

Brookes MJ, Woolrich M, Luckhoo H, Price D, Hale JR, Stephenson MC, Barnes GR, Smith SM, Morris PG (2011) Investigating the electrophysiological basis of resting state networks using magnetoencephalography. Proceedings of the National Academy of Sciences 108:16783–16788.

Brunet D, Murray MM, Michel CM (2011) Spatiotemporal analysis of multichannel EEG: CARTOOL. Computational intelligence and neuroscience 2011:2.

Cabral J, Vidaurre D, Marques P, Magalhães R, Moreira PS, Soares JM, Deco G, Sousa N, Kringelbach ML (2017) Cognitive performance in healthy older adults relates to spontaneous switching between states of functional connectivity during rest. Scientific reports 7:1–13.

Colclough GL, Brookes MJ, Smith SM, Woolrich MW (2015) A symmetric multivariate leakage correction for MEG connectomes. Neuroimage 117:439–448.

Contreras D, Steriade M (1996) Synchronization of low-frequency rhythms in corticothalamic networks. Neuroscience 76:11–24.

Deco G, Kringelbach ML (2016) Metastability and coherence: extending the communication through coherence hypothesis using a whole-brain computational perspective. Trends in neurosciences 39:125–135.

Deco G, Tononi G, Boly M, Kringelbach ML (2015) Rethinking segregation and integration: contributions of whole-brain modelling. Nature Reviews Neuroscience 16:430–439.

Destexhe A, Contreras D, Steriade M (1999) Spatiotemporal analysis of local field potentials and unit discharges in cat cerebral cortex during natural wake and sleep states. Journal of Neuroscience 19:4595–4608.

Engel AK, König P, Kreiter AK, Singer W (1991) Interhemispheric synchronization of oscillatory neuronal responses in cat visual cortex. Science:1177–1179.

Engel AK, Gerloff C, Hilgetag CC, Nolte G (2013) Intrinsic coupling modes: multiscale interactions in ongoing brain activity. Neuron 80:867–886.

Fox KC, Foster BL, Kucyi A, Daitch AL, Parvizi J (2018) Intracranial electrophysiology of the human default network. Trends in cognitive sciences 22:307–324.

Fries P (2005) A mechanism for cognitive dynamics: neuronal communication through neuronal coherence. Trends in cognitive sciences 9:474–480.

Ghuman AS, McDaniel JR, Martin A (2011) A wavelet-based method for measuring the oscillatory dynamics of resting-state functional connectivity in MEG. Neuroimage 56:69–77.

Gollo LL, Mirasso C, Sporns O, Breakspear M (2014) Mechanisms of zero-lag synchronization in cortical motifs. PLoS computational biology 10:e1003548.

Grave de Peralta Menendez R, Murray MM, Michel CM, Martuzzi R, Gonzalez Andino SL (2004) Electrical neuroimaging based on biophysical constraints. NeuroImage 21:527–539.

Greicius MD, Krasnow B, Reiss AL, Menon V (2003) Functional connectivity in the resting brain: a network analysis of the default mode hypothesis. Proceedings of the National Academy of Sciences 100:253–258.

He B, Sohrabpour A, Brown E, Liu Z (2018) Electrophysiological Source Imaging: A Noninvasive Window to Brain Dynamics. Annual Review of Biomedical Engineering 20:171–196.

He B, Astolfi L, Valdes-Sosa PA, Marinazzo D, Palva S, Benar CG, Michel CM, Koenig T (2019) Electrophysiological Brain Connectivity: Theory and Implementation. IEEE Transactions on Biomedical Engineering.

Hipp JF, Engel AK, Siegel M (2011) Oscillatory synchronization in large-scale cortical networks predicts perception. Neuron 69:387–396.

Hipp JF, Hawellek DJ, Corbetta M, Siegel M, Engel AK (2012) Large-scale cortical correlation structure of spontaneous oscillatory activity. Nature neuroscience 15:884.

Ju H, Bassett DS (2020) Dynamic representations in networked neural systems. Nature Neuroscience:1–10.

Jung T-P, Makeig S, Humphries C, Lee T-W, Mckeown MJ, Iragui V, Sejnowski TJ (2000) Removing electroencephalographic artifacts by blind source separation. Psychophysiology 37:163–178.

Krishnaswamy P, Obregon-Henao G, Ahveninen J, Khan S, Babadi B, Iglesias JE, Hämäläinen MS, Purdon PL (2017) Sparsity enables estimation of both subcortical and cortical activity from MEG and EEG. Proceedings of the National Academy of Sciences 114:E10465–E10474.

Lachaux J-P, Lutz A, Rudrauf D, Cosmelli D, Le Van Quyen M, Martinerie J, Varela F (2002) Estimating the time-course of coherence between single-trial brain signals: an introduction to wavelet coherence. Neurophysiologie Clinique/Clinical Neurophysiology 32:157–174.

Lachaux JP, Rodriguez E, Martinerie J, Varela FJ (1999) Measuring phase synchrony in brain signals. Human brain mapping 8:194–208.

Marzetti L, Della Penna S, Snyder AZ, Pizzella V, Nolte G, de Pasquale F, Romani GL, Corbetta M (2013) Frequency specific interactions of MEG resting state activity within and across brain networks as revealed by the multivariate interaction measure. Neuroimage 79:172–183.

Michel CM, Murray MM (2012) Towards the utilization of EEG as a brain imaging tool. Neuroimage 61:371–385.

Michel CM, Koenig T (2018) EEG microstates as a tool for studying the temporal dynamics of whole-brain neuronal networks: a review. Neuroimage 180:577–593.

Michel CM, Brunet D (2019) EEG Source Imaging: a practical review of the analysis steps. Frontiers in neurology 10.

Michel CM, Murray MM, Lantz G, Gonzalez S, Spinelli L, Grave de Peralta R (2004) EEG source imaging. Clin Neurophysiol 115:2195–2222.

Miller KJ, Weaver KE, Ojemann JG (2009) Direct electrophysiological measurement of human default network areas. Proceedings of the National Academy of Sciences 106:12174–12177.

Morlet J, Arens G, Fourgeau E, Giard D (1982) Wave propagation and sampling theory—Part II: Sampling theory and complex waves. Geophysics 47:222–236.

Nolte G, Bai O, Wheaton L, Mari Z, Vorbach S, Hallett M (2004) Identifying true brain interaction from EEG data using the imaginary part of coherency. Clinical neurophysiology 115:2292–2307.

O’reilly C, Elsabbagh M (2020) Intracranial recordings reveal ubiquitous in-phase and in-antiphase functional connectivity between homologous brain regions in humans. BioRxiv.

Palva JM, Wang SH, Palva S, Zhigalov A, Monto S, Brookes MJ, Schoffelen J-M, Jerbi K (2018) Ghost interactions in MEG/EEG source space: A note of caution on inter-areal coupling measures. Neuroimage 173:632–643.

Palva S, Palva JM (2012) Discovering oscillatory interaction networks with M/EEG: challenges and breakthroughs. Trends in cognitive sciences 16:219–230.

Raichle ME (2010) Two views of brain function. Trends in Cognitive Sciences 14:180–190.

Raichle ME, MacLeod AM, Snyder AZ, Powers WJ, Gusnard DA, Shulman GL (2001) A default mode of brain function. Proceedings of the National Academy of Sciences 98:676–682.

Roelfsema PR, Engel AK, König P, Singer W (1997) Visuomotor integration is associated with zero time-lag synchronization among cortical areas. Nature 385:157.

Rubega M, Carboni M, Seeber M, Pascucci D, Tourbier S, Toscano G, Van Mierlo P, Hagmann P, Plomp G, Vulliemoz S, Michel CM (2018) Estimating EEG Source Dipole Orientation Based on Singular-value Decomposition for Connectivity Analysis. Brain Topography 32:704–719.

Samogin J, Liu Q, Marino M, Wenderoth N, Mantini D (2019) Shared and connection-specific intrinsic interactions in the default mode network. Neuroimage 200:474–481.

Schaefer A, Kong R, Gordon EM, Laumann TO, Zuo X-N, Holmes AJ, Eickhoff SB, Yeo BT (2017) Local-global parcellation of the human cerebral cortex from intrinsic functional connectivity MRI. Cerebral Cortex 28:3095–3114.

Seeber M, Cantonas LM, Hoevels M, Sesia T, Visser-Vandewalle V, Michel CM (2019) Subcortical electrophysiological activity is detectable with high-density EEG source imaging. Nat Commun 10:753.

Seeley WW, Menon V, Schatzberg AF, Keller J, Glover GH, Kenna H, Reiss AL, Greicius MD (2007) Dissociable Intrinsic Connectivity Networks for Salience Processing and Executive Control. Journal of Neuroscience 27:2349–2356.

Siegel M, Donner TH, Oostenveld R, Fries P, Engel AK (2008) Neuronal synchronization along the dorsal visual pathway reflects the focus of spatial attention. Neuron 60:709–719.

Singer W (1999) Neuronal synchrony: a versatile code for the definition of relations? Neuron 24:49–65.

Smith SM, Fox PT, Miller KL, Glahn DC, Fox PM, Mackay CE, Filippini N, Watkins KE, Toro R, Laird AR (2009) Correspondence of the brain’s functional architecture during activation and rest. Proceedings of the National Academy of Sciences 106:13040–13045.

Stam CJ, Nolte G, Daffertshofer A (2007) Phase lag index: assessment of functional connectivity from multi channel EEG and MEG with diminished bias from common sources. Human brain mapping 28:1178–1193.

Tagliazucchi E, Balenzuela P, Fraiman D, Chialvo DR (2012) Criticality in Large-Scale Brain fMRI Dynamics Unveiled by a Novel Point Process Analysis. Frontiers in Physiology 3.

Tognoli E, Kelso JS (2014) The metastable brain. Neuron 81:35–48.

Tononi G, Edelman GM (1998) Consciousness and complexity. science 282:1846–1851.

Van Kerkoerle T, Self MW, Dagnino B, Gariel-Mathis M-A, Poort J, Van Der Togt C, Roelfsema PR (2014) Alpha and gamma oscillations characterize feedback and feedforward processing in monkey visual cortex. Proceedings of the National Academy of Sciences 111:14332–14341.

Varela F, Lachaux J-P, Rodriguez E, Martinerie J (2001) The brainweb: phase synchronization and large-scale integration. Nature reviews neuroscience 2:229.

Vicente R, Gollo LL, Mirasso CR, Fischer I, Pipa G (2008) Dynamical relaying can yield zero time lag neuronal synchrony despite long conduction delays. Proceedings of the National Academy of Sciences 105:17157–17162.

Vidaurre D, Hunt LT, Quinn AJ, Hunt BA, Brookes MJ, Nobre AC, Woolrich MW (2018) Spontaneous cortical activity transiently organises into frequency specific phase-coupling networks. Nat Commun 9:2987.

Von Stein A, Chiang C, König P (2000) Top-down processing mediated by interareal synchronization. Proceedings of the National Academy of Sciences 97:14748–14753.

Womelsdorf T, Schoffelen J-M, Oostenveld R, Singer W, Desimone R, Engel AK, Fries P (2007) Modulation of neuronal interactions through neuronal synchronization. science 316:1609–1612.

